# Relaxed selection can speed the evolution of complex adaptations

**DOI:** 10.1101/2024.07.09.602773

**Authors:** Jeremy Draghi, C. Brandon Ogbunugafor, Luis Zaman, Todd L. Parsons

## Abstract

Natural selection drives adaptive evolution and removes deleterious mutations; these effects are countervailing when a complex adaptation requires mutations that are initially deleterious when they arise, but beneficial in combination. While many models of this dynamic consider how genetic drift or other influences can aid valley crossing by weakening selection, we lack a general, analytical treatment of when relaxed selection might speed this type of adaptation. Here we use simulation and analysis to show that relaxed selection is generally favorable for valley-crossing when adaptive pathways require more than a single deleterious step. We also demonstrate that spatial heterogeneity in selection pressures could, by relaxing selection, allow populations to cross valleys much more rapidly than expected. These results relate to several applications of evolutionary theory to complex systems ranging from host-pathogen evolution to search algorithms in computer science.

## Introduction

Ever since Sewall Wright introduced the fitness landscape in 1931, the metaphor of mountains as genetic regions of high fitness separated by valleys of low fitness has been the canonical framework for understanding how epistatic (i.e., non-additive) interactions between genes impact evolutionary dynamics (1; 2; 3). From the beginning this framework has been applied to the origins of complex traits—those that seemingly require multiple, well-tuned parts to deliver adaptive benefits. Using the fitness landscape metaphor, adaptations in which several individually deleterious mutations improve fitness when combined are visualized by peaks separated by valleys. Quantifying the speed of valley-crossing may shed light on the origins of complex adaptations throughout the history of life (4), but has also been applied to contemporary problems like evolution in somatic cancers (5; 6). These same questions are also essential for predicting evolution in several applications that rely on the assumption that valley-crossing will be difficult: the use of multiple simultaneous treatments against bacterial infections (7; 8) and chronic viral infections like HIV (9), and the use of multiple resistance genes against plant pathogens (10).

Though valley crossing can in principle occur via the sequential fixation of each deleterious mutation, this pathway is a virtual impossibility in large populations when intermediates steps are significantly deleterious (11). Wright’s shifting balance theory avoided this problem by focusing on small populations which were hypothesized to occasionally shift to a different peak with the aid of genetic drift (3). The relevance of this model for evolution has been vigorously debated (see (12)), but more recent theory has shown that large populations can readily cross valleys and do so without substantial drops in mean fitness. Rather than sequential fixation, a lineage carrying a deleterious mutation can acquire subsequent mutations before the deleterious intermediates become very common (11; 5). This process may be slow, on average, but not gradual: distributions of crossing times are approximately exponential (13), reflecting the waiting time for a series of events that is rare but proceeds quickly once initiated.

Selection plays a unique, dual role in crossing adaptive valleys, manifesting as both the constraining and the driving forces. Consequently, it is not at all obvious whether more intense selection would speed or slow the process, and several models suggest that crossing is faster when selection is weakened or opposed. In Wright’s models, genetic drift played this countervailing role. Other candidate mechanisms that enable valley crossing by lessening the influence of natural selection include random phenotypic variation (14; 15), spatial (16; 17) or temporal (18; 19) heterogeneity in the strength of selection, high genetic variation for fitness (20; 21), non-Mendelian transmission biases (22), low density (23), and subdivided populations with local competition (24) or cooperative social interactions (25; 26). But despite these many examples and detailed mathematical analyses of some aspects of valley crossing, we still lack clear criteria for when lessened or relaxed selection speeds the arrival of complex adaptations.

Here we use analytical methods and simulations to show that relaxed selec-tion can speed valley-crossing when multiple deleterious steps separate peaks. We then examine the effects of one generic cause of relaxed selection in nature— spatial heterogeneity, in which populations spread across multiple environments experience relaxed selection on environment-specific traits. We model this scenario and show that populations faced with multiple, independent valley-crossing problems can surmount all of them in parallel, much more readily than would be extrapolated from the behavior of a specialist population in a single environment.

## Model & Results

### I. Valley crossing and relaxed selection

We define adaptive valleys following the terminology of (27). As shown in Figure 1, *k* mutations are required to realize a fitness benefit *s* relative to the wild-type. Each mutation individually lowers fitness by δ outside of the context of the *k* - mutant genotype (Figure 1). For simplicity, we follow (27) in modeling additive effects, such that the fitness *W*_*i*_ for 0 *<i< k* is 1 *<δi*, not (1 δ)^*i*^. This choice is an excellent approximation for the multiplicative case if δ and *k* are both small.

**Figure 1:**
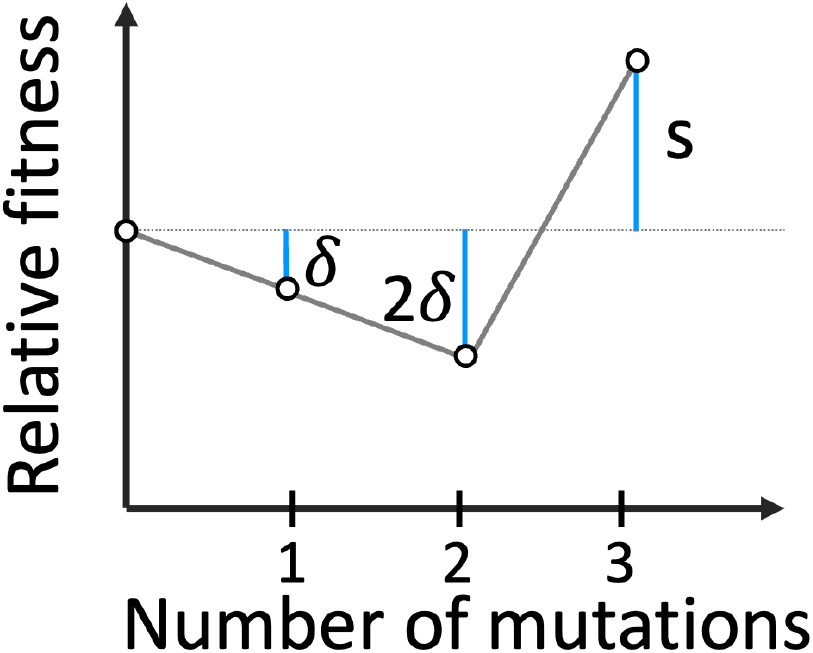
Illustration of the basic fitness landscape for the *k* = 3 case.

We model asexual reproduction in haploids. Each offspring carries a Poisson-distributed number of mutations with mean *µ*, and the total mutations carried by a genotype is capped at *k*. For simplicity, back-mutation is not allowed, and the forward mutation rate does not depend on the number of mutations carried by the parent.

Waiting times for valley-crossing have been analyzed systematically across a number of papers (e.g., (28; 27)). However, almost all such results focus on two-step crossings (*k* = 2) in the case where *<* is substantial relative to the strength of drift, or instead consider more than two steps but with effectively neutral intermediates (29; 30; 31). A few studies formally consider cases where *k>* 2 for deleterious intermediates, but have not analyzed the consequences of relaxed selection for crossing time (27; 32; 33). In the supplemental material, we analyze waiting times for arbitrary *k* for both the Moran and Wright-Fisher models. Below we sketch our main results and compare the resulting predictions to simulations.

First, as a point of comparison we examine the effects of relaxed selection on *k* = 2 valleys. For a Moran process, we can approximate the waiting time until a genotype with *k* mutations arises as the product of the number of first-step mutations per generation, *Nµ*, the expected lineage size created by each such mutation, δ^−1^, and the mutation rate from one to two mutations, *µ*. This calculation assumes that deleterious steps are truly deleterious (δ ≫*N*^−1^) and mutations relatively rare (*µ ≪*1), ensuring that the first-step mutation remains rare. Once the second mutant arises, its chance to fix is well-approximated as 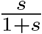 for large *N*. Relaxed selection, achieved by multiplying both *δ* and *s* by the same constant, is therefore expected to have little effect on the arrival time of the k-mutant for *k* = 2 as this constant will largely cancel out of the expected waiting time when *s ≪* 1 + *s*. Further, the expected waiting time until the *k*-mutant has achieved a substantial frequency will only increase with weaker selection. In short, we do not expect relaxed selection to have a large or consistent effect on mean crossing times when *k* = 2.

For larger valleys (*k>* 2), these dynamics are qualitatively different. Figure 2 depicts landscapes of the probability of crossing valleys of for *k* = 2, 3, and 4. For *k* = 2, weakening selection (movement along the diagonal lines toward the lower-left corner) has little effect on this probability, as the contour lines are nearly parallel to this vector of movement. But for higher *k*, there are large regions of parameter space in which weakening selection substantially raises the probability of crossing valleys.

**Figure 2:**
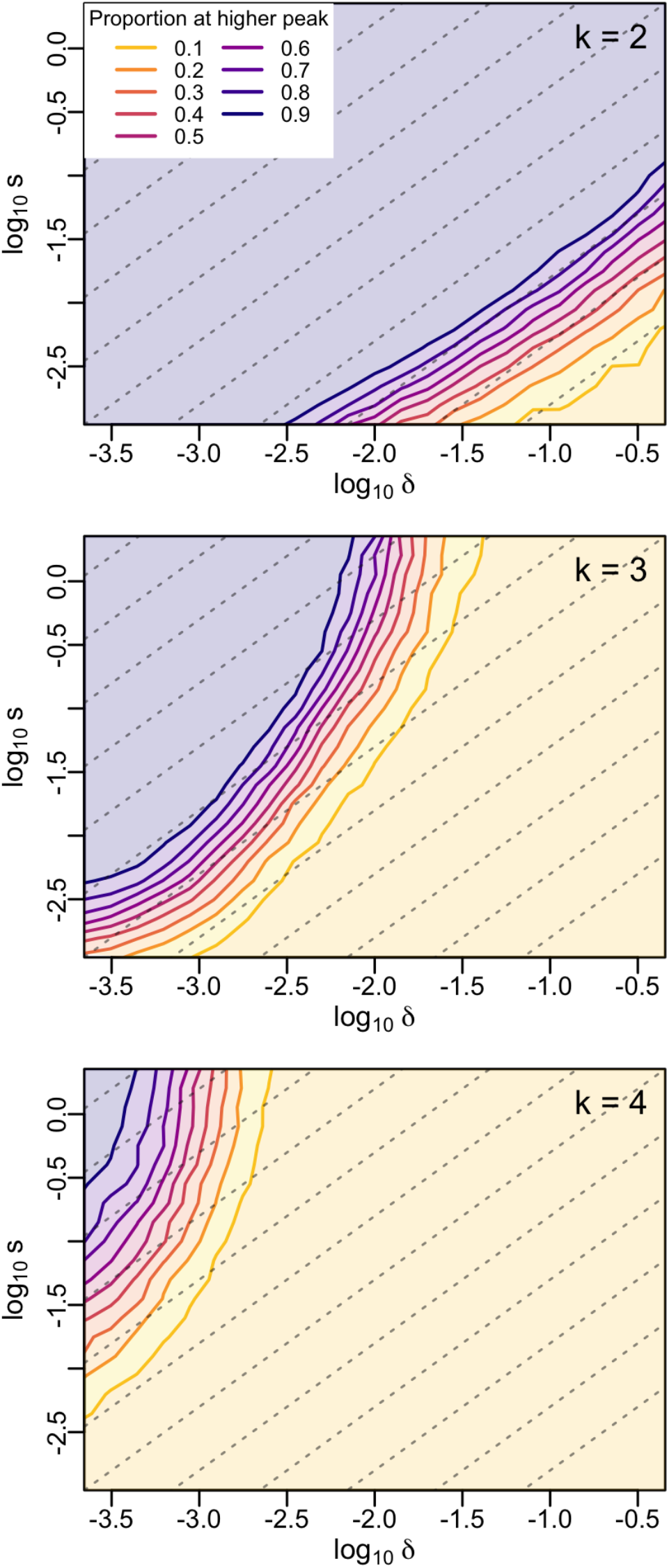
Probability of the *k*-mutant to arise and invade (reach a frequency of at least 0.01) within 5000 generations for three values of *k*. The Wright-Fisher model was simulated with *N* = 500, 000 and *µ* = 0.00005 across more than three orders of magnitude for both *δ* and *s*. B. Diagonal lines show proportional change in both *δ* and *s*; movement along these lines from the top-right to the bottom-left corresponds to weakening selection. A minimum of two hundred replicate simulations were performed for each point in a 21 × 21 grid of conditions for each value of *k*.

We analyzed these dynamics in both the Moran and Wright-Fisher models, deriving approximations for the arrival time of the first mutant that will be ancestral to a successful *k*-mutant lineage (see Supplemental Materials §1.2). Under the assumption that the lineages seeded by deleterious intermediates are expected to remain small 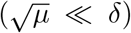 and that the k-mutant is strongly favored, we can generate accurate, simple formula for the rate at which valleys are crossed. When 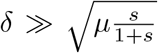, we derive a particularly intuitive expression for the *k* = 3 waiting time in the Moran model (see Supplemental Materials, Equations S.14–16):

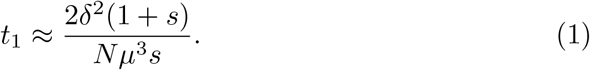

This expression can be decomposed into the flux of first-step mutations per generation (*Nµ*), the number of second-step mutations they give rise to 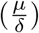, the number of *k*-mutants arising from each of those 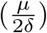, and the fixation probability of each of those *k*-mutants 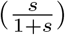. Together, this expression illustrates why relaxed selection speeds valley-crossing for larger valleys: because there are multiple deleterious steps, adaption is accelerated if *δ* is reduced, even is *s* is reduced by the same amount. In general, we find a dependency of order *δ*^*k-*1^, indicating that the benefits of relaxed selection increase with *k* (see Supplemental Materials §1.2.2).

When *s* is relatively large compared to *µ*, we expect that the vast majority of the waiting time for a successful *k*-mutant will be spent waiting for the first-step ancestor described by Eq. 1. Therefore, we can take *t*_1_ as a good approximation of *t*_*k*_, the arrival time of the *k*-mutant (see Supplementary Materials §4). Figure 3 plots simulated observations of these arrival times, along with time, *t*_inv_, at which the k-mutant has reached a frequency of 0.1. Relaxing selection, modeled as multiplying both *s* and *δ* by the fraction indicated on the x-axis, substantially lowers the waiting time. For the Moran model, eq. 1 accurately captures this trend as long as 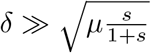, visualized as the transition from a solid to a dotted line. When selection is very weak, *t*_1_ and *t*_inv_ separate noticeably, mitigating any further effect of decreasing selection on valley-crossing times. In the Supplemental Materials, we derive similar formula for the Wright-Fisher model (see Supplementary Materials §4). Moreover, we show that relaxed selection is generally expected to speed the arrival of the *k*-mutant as long as *k >* 2 (see Supplemental Materials §1.3).

**Figure 3:**
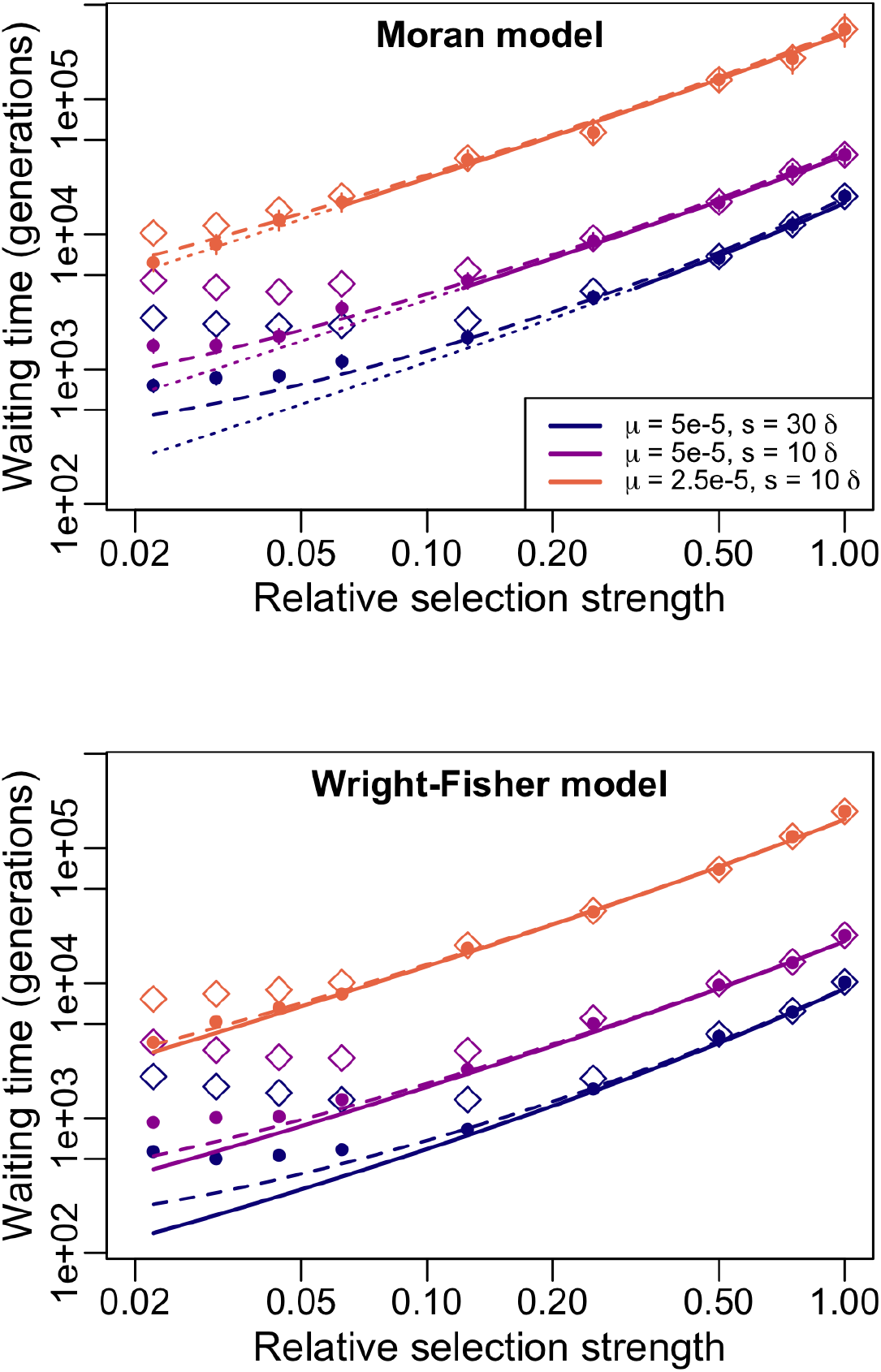
Mean waiting times of valley-crossing for *k* = 3 in the Moran (*top*) and Wright-Fisher (*bottom*) models. Filled points show the means of *t*_1_ from replicate simulations; open diamonds show the corresponding mean of *t*_inv_. (Top) Solid lines indicate predictions from Eq. 1; the line is dotted when the ratio of *δ* to 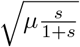 is below five, indicating that we expect the approximation to begin to fail. Dashed lines are calculated iteratively from Supplemental Materials Eq. S.5. (Bottom) Solid lines indicate predictions from the approximations given by Eq. S.40; dashed lines show the more complete expression given by iterating Eq. S.34. Confidence intervals, determined by bootstrapping, are visible where they exceed the size of plotting symbols.

### II. Valley crossing and spatial heterogeneity

Relaxed selection can arise when a population is spread over multiple environments, and a trait, or the genes underlying it, are not under selection in some of those environments. To explore how this form of relaxed selection could impact valley crossing, we simulated populations distributed among *m* environments. In each environment, we assumed that fitness was determined by a set of loci expressed uniquely in that one environment. Alleles expressed within each environment were subject to a valley-crossing scenario, exactly as described above. For example, with *m* = 5 and *k* = 3 a population would be spread across five environments at any one time and faced with five genetically distinct valley-crossing problems, represented by a genome of fifteen total loci. Each valley-crossing dynamic would then be allowed to evolve via separate mutations, with the fitness of an individual being determined by these mutations and their environment.

Within this basic scenario of spatial heterogeneity, competition and selection can operate in several different ways. An individual organism may experience only one environment within their lifespan (coarse-grained heterogeneity) or may visit many environments (fine-grained heterogeneity). In the former case we uniformly assigned each individual at birth to one of the *m* environments, eventually filling each environment with *N/m* organisms. In the latter, we assigned a fitness to each individual by averaging its fitness across all *m* habitats. Separately, selection can occur by global competition (also called hard selection) or can be localized to each environment (soft selection). To simulate the former, we chose *N* individuals for the next generation in proportion to assigned fitnesses; to simulate soft selection, we first standardized fitnesses within each environment by dividing them by the mean in that environment. We therefore model four different combinations of these decisions, which represent extremes on spectra bridging coarse- and fine-grained heterogeneity and global and local selection.

Figure 4 illustrates that populations can cross valleys much more readily when spread across multiple environments. While local competition is associated with slightly more facile valley-crossing, the main effect of spatial heterogeneity is largely independent of the details of selection. Note that these probabilities are averaged across each environment, such that the number of valleys crossed is the product of *p*_inv_ and the number of environments. For example, a population spread across ten environments typically crosses more than four valleys; in the same span of time, a population experiencing only a single environment would have about a 10% chance to cross that single valley.

**Figure 4:**
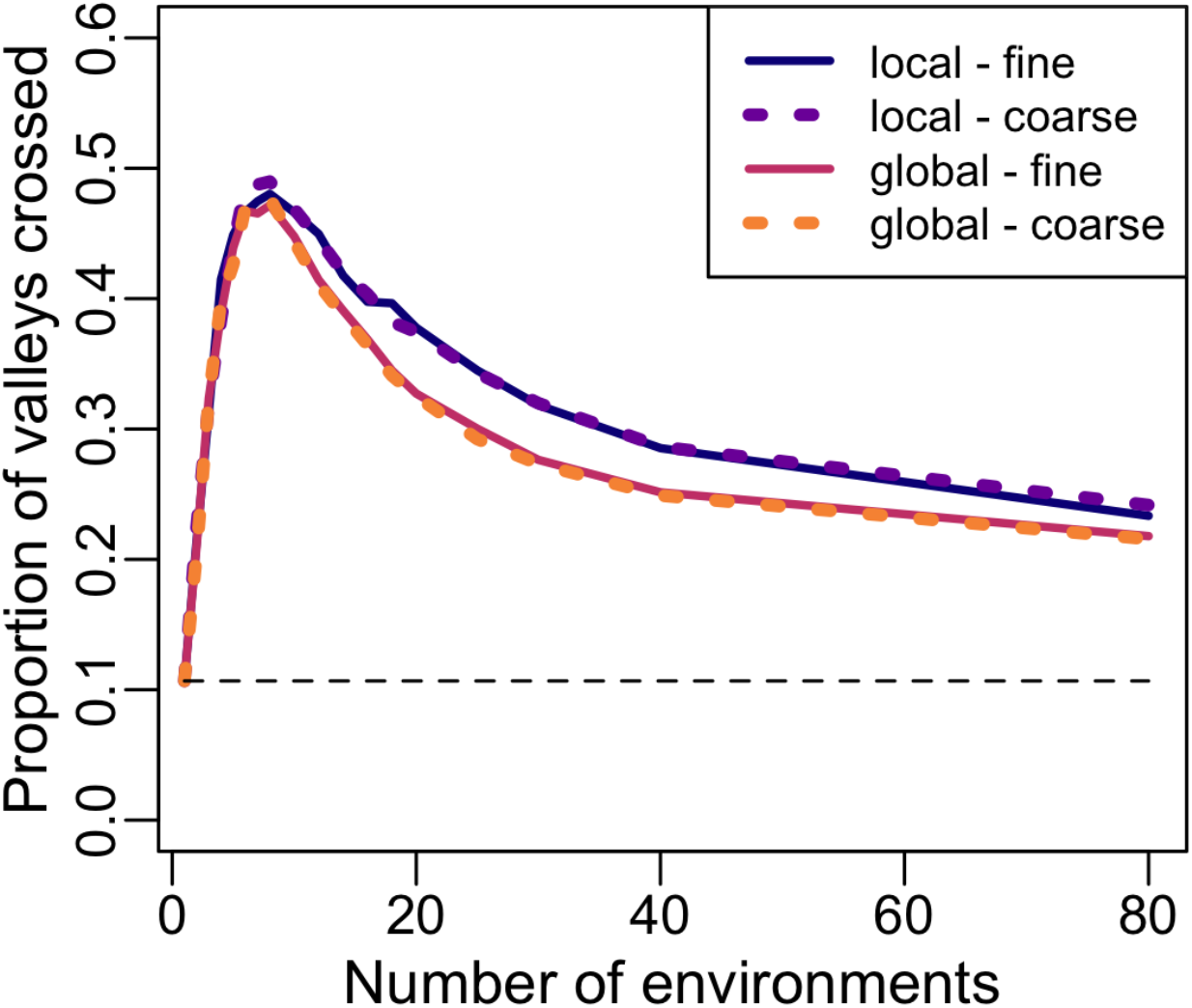
Probability of the *k* = 3 mutant to arise and invade (reach a frequency of at least 0.01) within 5000 generations, averaged over each environment the population experiences (x-axis). Each population is spread across *m* environments, with three loci per environment to create an independent *k* = 3 valley for each environment. The Wright-Fisher model was simulated with *N* = 504000 and *µ* = 0.00005 (the population size was chosen so that it was divisible by each tested value of *m*). The dashed line shows the expected probability for a single environment, as determined by simulation. At least three hundred replicatesare averaged for each treatment.

## Discussion

Empirical fitness landscapes motivate the need for quantitative predictions of how populations with specific characteristics, and in particular environments, can navigate their ways to higher fitness (34; 35; 2). Strictly uphill paths are often evident in cases of evolution under strong, focused sources of selection (36; 37). In other cases, valleys may be present but can be skirted by evolution along ridges (1): mutations comprising the complex adaptation can be acquired in sequences of substitutions, or in particular environments, in which each step is individually beneficial, following paths that are lengthy but still entirely uphill (e.g., (38; 39)). More generally, though, fitness landscapes may have properties that make evolutionary optimization a *hard* problem in computational terms (40). In practical applications of valley-crossing theory like the acquisition of multi-drug resistance in HIV (19), we must study the causes of evolutionary events that are rare precisely because we have sought to design treatments that are hard for evolution to undermine. The example of resistance to treatment cocktails in HIV illustrates that understanding valley crossing is particularly important in applied contexts in which we have constructed situations that frustrate simple adaptation. In this study, we look at two general approaches to slowing adaptation—adaptive valleys and relaxed selection caused by spatial heterogeneity. Surprisingly, these two obstacles to adaptive evolution interact antagonistically, partially canceling each other’s constraining effect when certain conditions are met. When adaptive valleys are wider than the minimum size of *k* = 2, and organisms experience orthogonal valley-crossing problems in distinct environments, then for a broad range of parameters environmental heterogeneity speeds up valley crossing.

Virus evolution, in particular, has a long history of applications of theoretical population genetics to adaptation on rugged fitness landscapes like those modeled in this study. This is partly because viruses embody the properties (e.g.,large population sizes and high mutation rates) that facilitate rapid adaptation to diverse environments(41). More recently, the origin of the SARS-CoV-2 variants has drawn attention to the relevance of rapid evolution across epistatic landscapes(42). For example, the SARS-CoV-2 Omicron (B.1.1.529) variant contains several mutations that are likely to be deleterious individually but beneficial in combination (43). If, as is hypothesized, Omicron originated in chronic infection in an immunocompromised host, this environment likely facilitated the fixation of mutations (corresponding to immune escape and/or increased transmission) that were otherwise too deleterious individually to permit valley crossing in immunocompetent individuals (44; 45); indeed, relaxed selection is explicitly hypothesized as an important driver of Omicron’s emergence (46).

Similar ideas apply to the use of vaccines, which may owe some of their therapeutic longevity to the varied antibodies they induced across the host population, creating a spatially heterogeneous set of selective pressures for pathogens (47). Experimental evidence in prokaryotes demonstrates the importance of a diversity of immune responses (48) and models have analyzed the benefits of distributing varied vaccines for fighting infections (49). Yet, our results suggest that presenting pathogens with a varied set of evolutionary challenges may not be as effective a constraint on escape as one might hope. In SARS-CoV-2, different components of viral fitness show strong epistasis among mutations that might facilitate escape (50; 46; 51). Moreover, host antibodies targeting one virus epitope have limited effects on binding to other epitopes, suggesting a heterogeneous selective environment for virus mutants (51). Our results suggest that while epistasis, and the resulting fitness valleys, may constrain viral evolution, the fact that diverse virus epitopes are targeted by distinct antibodies across hosts may weaken, rather than strengthen, this constraint.

Complimenting our results, recent work modeling pathogen populations (23) has noted that a type of relaxed selection can arise from low contact rates between infected individuals, allowing deleterious mutants to persist without being immediately out-competed by coinfecting wild-type strains, consequently facilitating valley crossing. Future reconciliations between evolutionary theory and the dynamics of infectious diseases could more clearly identify epidemiological scenarios that promote or prevent the emergence of variants by identifying the population genetic conditions where valley crossing and immune escape (via adaptation on rugged fitness landscapes) is more likely.

While our findings are relevant to real world problems in infectious diseases, the implications are broader. Valley-crossing in evolution is formally very similar to the problem of trapping at local minima in energy landscapes or premature convergence in evolutionary computing. In part, this can be understood as a dilemma of exploration vs exploitation: the greater the strength or stringency of natural selection, the more the evolutionary process is slanted toward exploiting already-discovered high-fitness genotypes, rather than exploring the potential offspring of genotypes with lower fitnesses in the present. In the context of directed evolution of macromolecules, researchers have connected the benefits of relaxed selection to the magnitude of epistasis (52), mirroring the epistasis central to the valley-crossing examples studied here.

The field of evolutionary computation in computer science has long examined analogous problems, where the search for optimal solutions in high-dimensional rugged parameter spaces led researchers to algorithmic implementations of Darwinian evolution. The problems being solved often require multi-objective selection, in which several sub-problems are simultaneously optimized and sophisticated algorithms are used to ensure progress is made along all fronts. Recently, these multi-objective algorithms have been tested for their efficacy in directed evolution experiments using microbial populations and show great promise (53; 54). The balance of adaptation being spread across multiple objectives, allowing each to afford some genetics drift on the other fronts, is a feature of these multiobjective approaches that we argue is contributing to their success. Other concepts rooted in computer science but with interdisciplinary applications include deception (55). Deceptive landscapes contain features that lead search processes away from global optima, as was recently shown to be the case for some antibiotic-resistance landscapes (56). These examples point to the largely unexplored potential for cross-disciplinary progress on best to navigate rugged landscapes.

Other surprising effects have been investigated using these more computational and engineering approaches, some of which can be attributable to the mechanism we investigate here. For example, Dolson and Ofria observed that evolutionary hostspots–regions of the physical world that produced a disproportionately large amount of novel variation–often occurred in regions of high resource overlap (57). As in our models, such diversity of adaptive opportunities could increase the effects of drift, and thus allow populations to cross more or larger fitness valleys. Recent computational work by Bohm et al. shows how transient relaxed selection, which is driven by the lagging increase in mean fitness of an adapting population, leads to temporary increases in exploration of the fitness landscape (21). Pushing the limit of relaxed selection further, *novelty search* is a new selection algorithm that has gained substantial popularity and success, where rather than rewarding any single (or multiple) objective, individuals are selected based on the uniqueness of their phenotype (58). It is counter-intuitive that jettisoning the evaluation of success actually improves the search for optimal solutions, but our results perhaps offer a simple explanation: relaxed selection allows valley crossing that would otherwise inhibit evolutionary exploration.

Understanding the dynamics of valley-crossing in adaptive landscapes has become a pragmatic, multi-disciplinary problem. Classically, fitness landscapes have helped depict evolution as a set of genotypes competing in a single environment. Our results here help to emphasize the importance of considering environmental variation: when populations are spread across environments, each with its own fitness landscape, our expectations for important dynamics like valley crossing are substantially different. While we have focused here on a simplified picture, without pleiotropy, of how environment-specific landscapes interact. Future work could continue to explore how interrelated landscapes across environments, in which mutations can have pleiotropic effect, shape the diversity of trajectories and likelihood of progress toward higher fitness peaks.

## Acknowledgements

This work was partially supported by NSF (USA) Division of Environmental Biology award 2147101 to JAD and TLP, the NSF Division of Environmental Biology Award Number 2142720 to C.B.O, and the NSF Division of Environmental Biology Award Number 1813069 to L.Z.

## Methods

Individual-based simulations were used to track evolution of distinct genotypes in populations of fixed size, with reproduction occurring with overlapping (Moran) or discrete (Wright-Fisher) generations. In both types of simulations, we tracked the lineages of all genotypes such that we could unambiguously es-tablish the origination time of each mutation in a successful lineage. Matching the assumptions of the analytical model, we assumed that the chance of a mutational change from *i* to *i* + 1 mutations was independent of *i*. However, unlike the assumptions of the analysis we allowed for the possibility of multiple mutations in a single reproductive event. Selection affected fecundity in both model. Simulations were coded in R and C.

## Data sharing plan

All code and simulation data will be made available on Data Dryad prior to publication.

## Supplementary Material

In what follows, we obtain estimates for the valley crossing time for a Moran model [13], allowing for a valley of arbitrary width (*i*.*e*., an arbitrary number of mutations, *k*, in which each is deleterious but the combination of all *k* is beneficial. We will focus on a model with ***fecundity selection*** (*i*.*e*., each epistatically negative mutation acquired reduces the individual’s birth rate), but proceeding identically, one can obtain a similar result for a model with ***mortality selection*** (*i*.*e*., when epistatically negative mutations increase an individual’s death rate).

After first specifying our model (§1), we present an approximation by a birth-and-death process with immigration [10] in §1.1. We then adapt a heuristic argument presented in [17] for the waiting time for *m* mutations in a neutral Moran model to our case (§1.2). While we content ourselves with an informal approach, these arguments can be rigorously justified using coupling arguments similar to those in [17, 6]. By considering various regimes, we are able further simplify our expressions (§1.2.1,1.2.2). Similar results appear in [19], obtained via a different and arguably more complicated argument. Using the analytical expression thus obtained, we show that relaxing selection reduces the expected valley-crossing time (§1.3).

We then illustrate the generality of our approach by applying it to the Wright-Fisher model (§2), obtaining what we believe to be the first estimates for that model. Interestingly, because we are working in a regime when the diffusion approximation is not appropriate (see *e*.*g*., [14] for a comparison of the exact Wright-Fisher model, its diffusion approximation, and an approximating branching process), we find that, unlike many quantities of population genetic interest, the expected time to cross a fitness valley is *not* identical in the two models, even after one accounts for the factor of two difference in the effective population sizes for the Moran and Wright-Fisher models [7, §3.7].

Finally, we provide evidence for the two key assumptions underlying our approach, that mutant individuals remain rare until the valley is crossed (§3), and that the time to valley crossing is dominated by the wait until the arrival of the first mutant individual that will be the ancestor of the adapted *k*-mutant that will go on to fix (§4).

We start from a general formulation, making simplifications as needed to obtain tractable analytical results. Equation (S.16) specializes to Equation [1] in the Main Text when one takes *k* = 3 and *i* = 1.

## 1 Moran Model

We assume that population grows and changes according to a Moran model in a finite population of size *N* [13]. The individuals are grouped into types *i* = 0,…, *k*, the type indicating the number of loci at which the individual carries a mutation, and we let *X*_*i*_(*t*) denote the number of individuals of type *i* at time *t*.

Selection can be modelled in the Moran framework in a number of asymptotically-equivalent ways; here we assume *fecundity selection*. Individuals of type *i* are assumed give birth at rate *δ*_*i*_. When a birth occurs, the offspring then displaces an individual chosen uniformly at random from the population (which, as formulated below, may include their parent). By assumption, individuals carrying 0 *< i < k* mutations are less fit than the wild-type, *δ*_0_ *> b*_1_ *>* …*> b*_*k-*1_, whereas an individual carrying *k* mutations enjoys a positive epistatic effect, and is more fit *δ*_*k*_ *> b*_0_.

If, in addition, we assume that the offspring of an individual with *i* mutations has *j* mutations with probability *µ*_*i,j*_ *≪*1 (*µ*_*j,j*_ = 0), then type *i* individuals are replaced by type *j* individuals at total rate

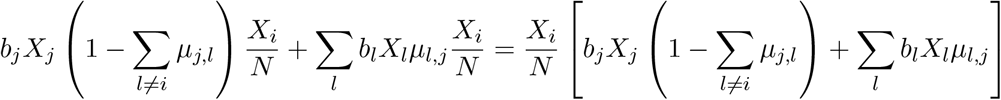

Thus, the number of type *i* individuals decreases by 1 at rate

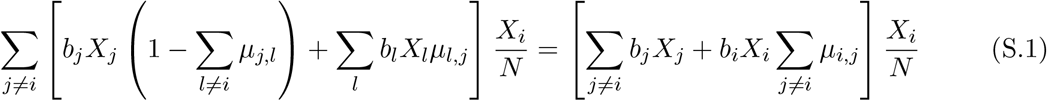

and increases by 1 at rate

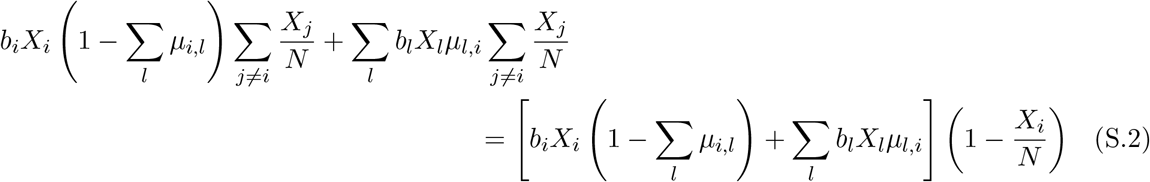

### 1.1 Birth-and-Death Process Approximation

By rescaling time, we can always take *δ*_0_ = 1, *δ*_*i*_ = 1 - *δ*_*i*_ for 0 *< δ*_1_ *<* …*< δ*_*k*−1_, and *δ*_*k*_ = 1 + *s* for *s >* 0. Moreover, suppose the number of *i* individuals is negligible compared to *N* whereas *X*_0_ ∼ *N*. This is assured by deleterious selection for individuals 0 *< i < k* mutations (see §3). In general, this is *not* the case for individuals with *k* mutations. However, we will only need the approximation up until the appearance of the first individual with *k* mutations that is destined to go on to fixation; up until this individual appears, *X*_*k*_ ≪ *N* (see §3.3).

Under these assumptions, for 0 *<i* (S.1) becomes

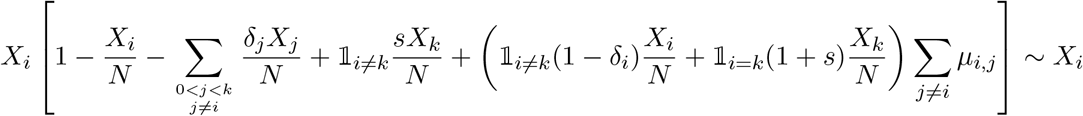

where 𝟙 _*i*≠*k*_ and 𝟙 _*i*=*k*_ are indicator functions (*i*.*e*., *𝟙* _*i*≠*k*_ is 1 if *i* ≠*k* and 0 if *i* = *k*, the opposite for 𝟙 _*i*=*k*_). For 0 *<i< k*, (S.2) becomes

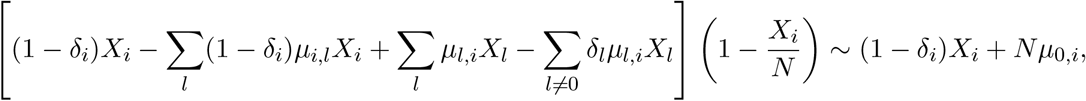

and, similarly, for *i* = *k*, (S.2) is approximately 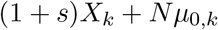.

Thus, if 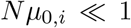_,*i*_ ≪ 1, we may approximate *X*_*i*_ by a simple birth-and-death process [10] with per-capita birth rate 1 − *δ*_*i*_ if *i<* 0 *< k*, and 1 + *s* if *i* = *k*, and per capita death rate 1. If, on the other hand, *µ*_0,*i*_ is of order 𝒪 (1), then we need to use a birth-and-death process with immigration [10] with the same birth and death rates and immigration rate 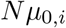.

### 1.2 sValley Crossing Rate

We adapt the heuristic presented in [17] to estimate the expected waiting time to cross a fitness valley. We do so by computing the probability that a given type 1 individual (*i*.*e*., one carrying the mutant allele at the first locus) is ***ancestral*** to a successful adapted individual. Here we define ***success*** by stipulating that the lineage of that individual must be destined to fix; a successful adapted individual has crossed the fitness valley. We will refer to this type 1 individual as ***ancestral*** or as the ***ancestor***. Below, we will argue that the time from the birth of the ancestor to the birth of the selected mutant destined to fix is negligible with respect the wait for the ancestor to appear. Thus, expected valley-crossing time, defined as the wait until the k-mutant appears, is approximately equal to the waiting time to the birth of this ancestor.

We henceforth assume that the probability of multiple mutations is negligible, so that an off-spring born to an individual carrying *i* mutations carries *i* + 1 mutations with probability *µ*_*i*_ and *i* mutations with probability 1-*µ*_*i*_, and no other mutations are possible (*i*.*e*., *µ*_*i,j*_ = *µ*_*i*_ if *j* = *i*+1, and *Nµ*_*i,j*_ ≪ 1 otherwise). We thus approximate the number of type 1 individuals by a birth-and-death process with immigration, and all other types *i>* 0 by birth-and-death processes.

As we observed above, prior to the selective sweep that follows the arrival of the mutant destined to fix, the population is almost entirely composed of wild type individuals, and using our approximations above, type 1 individuals are being produced at constant rate

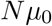

*i*.*e*., they are being produced at the times of a Poisson process with this rate, and the times between arrivals are exponentially distributed with rate *Nµ*_0_. Below, we will compute the probability, *p*_1_, that a given mutant is the ancestor. Then, using the thinning property of the Poisson process, an ancestor is produced at rate *Nµ*_0_*p*_1_, the waiting time until the it appears is exponentially distributed with rate *Nµ*_0_*p*_1_, and the mean time to the arrival of the ancestor is 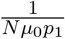.

#### 1.2.1 Ancestral Recursion

We compute *p*_1_ via recursion. As we saw above, we can approximate the Moran process by a birth-and-death process with type-dependent per-capita birth and death rates *δ*_*i*_ and *d*_*i*_:

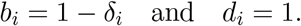

Let *p*_*i*_ be the probability that an individual carrying mutant alleles at *i* loci is ancestral to the successful individual, and consider the first event that happens to that individual during their lifetime (either a birth or death). The first event is a birth with probability

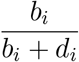

and a death with probability

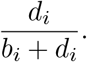

Clearly, if the first event is a death, the individual cannot be ancestral to a selected individual. If the first event is a birth, then the focal individual is ancestral if either it or its offspring is ancestral. Because our simple Markov model has no aging (*i*.*e*., in the event that the first event is a birth, the probabilities that the next event is a birth or a death are identical to those for the first event, *etc*.) the parent still has probability *p*_*i*_ of being ancestral after the first birth. On the other hand, there are two scenarios in which the offspring is ancestral: either it was born without mutation – with probability 1 − *µ*_*i*_ – and it has *i* mutations like its parent, and thus probability *p*_*i*_ of be ancestral, or, a mutation occurred – with probability *µ*_*i*_, and the offspring carries *i* + 1 mutant alleles, and thus has probability *p*_*i*+1_ of being ancestral. Combining these, the offspring has probability

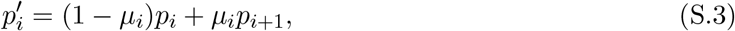

of being ancestral. Thus, the probability that at least one of the parent or the offspring is ancestral is

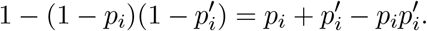

Combining all of the above, we see that

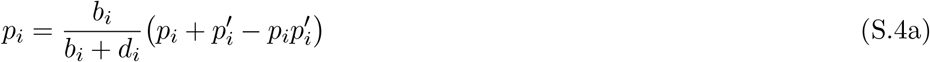

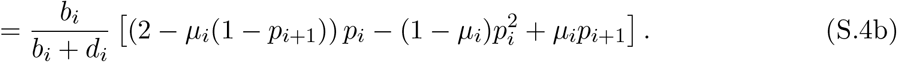

Rearranging, we have

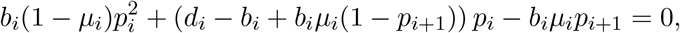

which may be solved via the quadratic equation to express *p*_*i*_ as a function of *p*_*i*+1_:

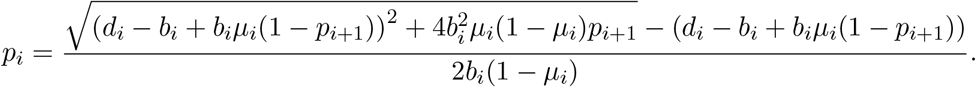

In particular, given an initial condition *p*_*k*_, we can iterate this to obtain *p*_1_.

Specializing to the case *δ*_*i*_ = 1 - *δ*_*i*_ and *d*_*i*_ = 1 gives us a slightly simpler expression,

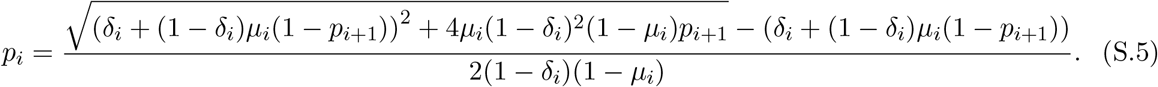

**p**_**k**_: Now, by assumption, individuals carrying *k* mutations are again adapted: they have birth rate *δ*_*k*_ = 1 + *s* and death rate *d*_*k*_ = 1. Then, using Moran’s expression for the fixation probability [13], the probability that an individual with *k* mutations is ancestral to the whole population is

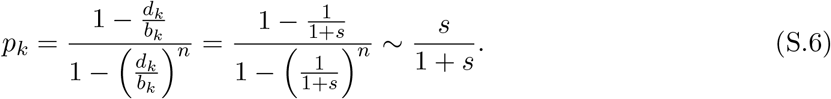

**p**_**k***-***1**_: Substituting (S.6) into (S.5) yields

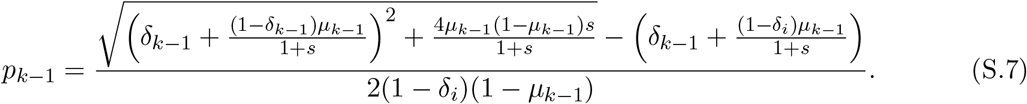

giving us a starting point for the recursion.

Unfortunately, the iteration quickly becomes unwieldy. If, however, we assume in addition that *µ*_*i*_ ≪ *δ*_*i*_ ≪ 1, then discarding all terms except those of highest order, (S.5) and (S.7) simplify to

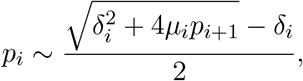

and

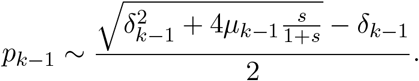

If, moreover, 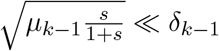, we have, using the binomial series 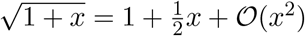,

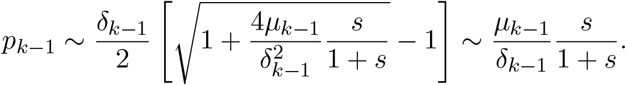

On the other hand, if 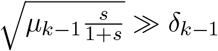, then

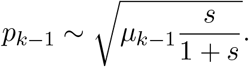

*N*.*δ*., when 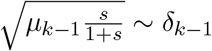, these last simplifications are not possible.

**p**_**k***-***2**_: Now, consider *p*_*k-*2_:

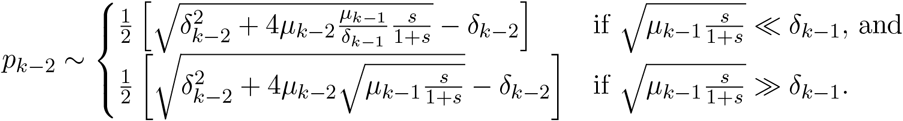

If we assume that all the *δ*_*i*_ have the same order of magnitude:

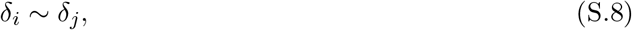

then 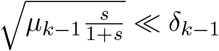 implies that 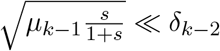 and

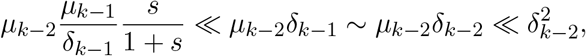

so the first case reduces, as above, to

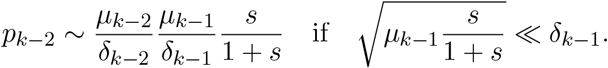

If, on the other hand, 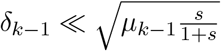, we need to consider two possibilities, either

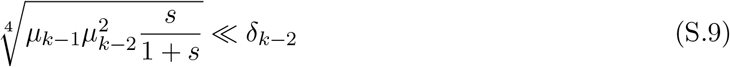

(*N*.*δ*., since, by assumption (S.8), *δ*_*k*− 2_ ∼ *δ*_*k*−1_, we also have 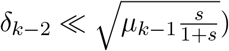 or,

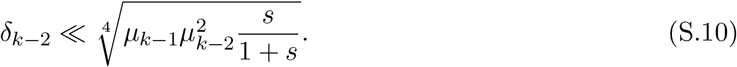

If we assume (S.9), then

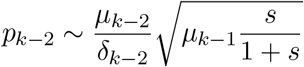

whereas assuming (S.10), we have

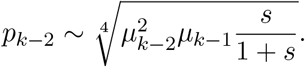

**p**_**k***-***l**_, **l** = **1**,…, **k** - **1:** Proceeding similarly, we find that if for some *i, i* = 1,…, *l* + 1

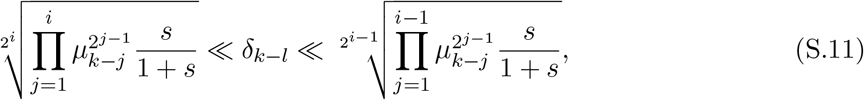

(*N*.*δ*.,, by our assumption (S.8), if this asymptotic inequality holds for any *δ*_*m*_, *m* = 1,…, *k* - 1, then it holds for all of them) then

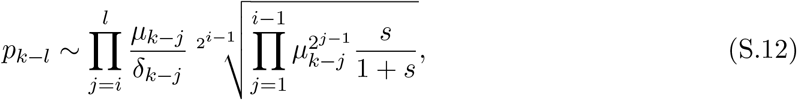

where we take 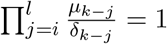 if *i> l*, and 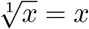. In particular, the waiting time until the arrival of the first adapted individual has mean

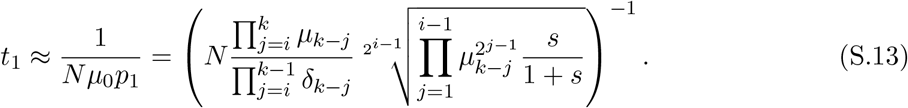

#### 1.2.2 Additive and Multiplicative Selection

Simplifying to additive (*δ*_*i*_ = *bi*) or multiplicative selection (1 + *δ*_*i*_ = (1 + *δ*)^*i*^, so *δ*_*i*_ ∼ *δi*, since, by assumption *δ*_*i*_ ≪ 1), and assuming that all mutation rates are equal (*µ*_*i*_ ≪ *µ*), we see that if

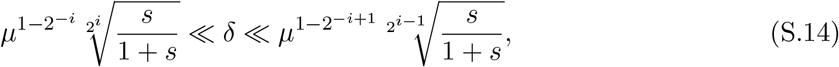

then, taking 0! = 1 and, as previously, 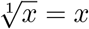,

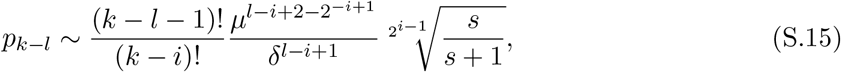

and

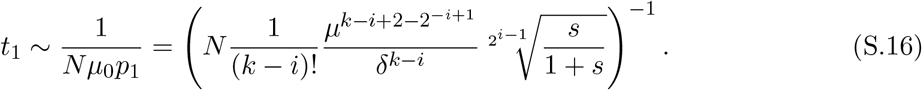

Equation [1] in the Main Text is obtained by taking *k* = 3 and *i* = 1.

### 1.3 Relaxing Selection Can Accelerate Adaptation

Now, suppose that all forms of selection are relaxed by a factor of *x* ∈ (0, 1), *i*.*e*., we replace *δ*_*i*_ by *δ*_*i*_*x* and *s* by *sx* above. This gives a probability of being ancestral that depends on *x*:

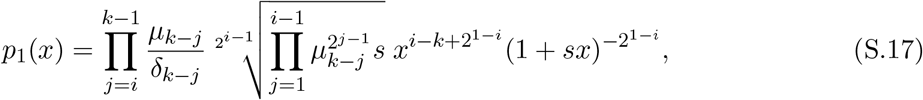

To see the effect on adaptation time, note that *increasing p*_1_ has the consequence of *decreasing* the mean adaptation time *t*_1_, (S.13). Now,

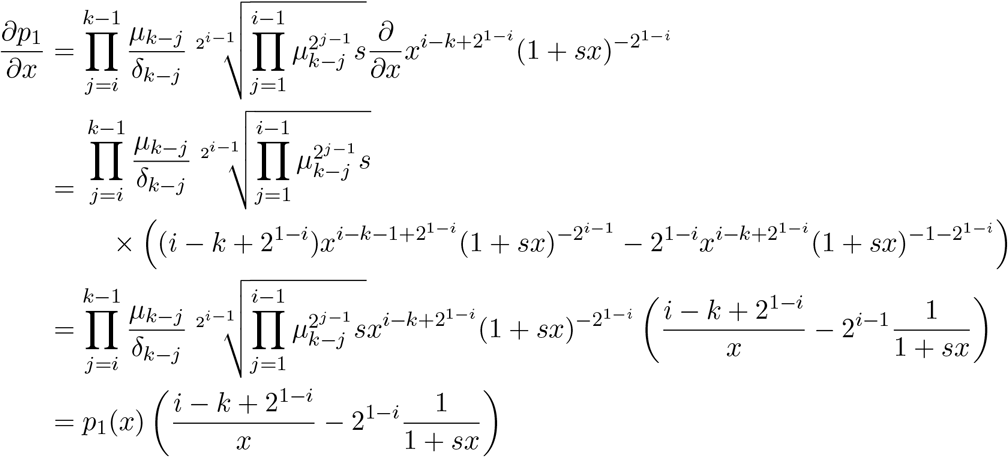

which is negative (so increasing with *decreasing x*) provided

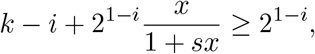

Now, for 0 *<x<* 1,

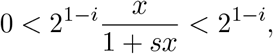

so the inequality holds provided *k* ≥ *i* + 1. Note that *k* ≥ *i*, whereas when *i* = *k*, the *δ*_*i*_ are in their smallest possible range (*cf*. (S.11)) and

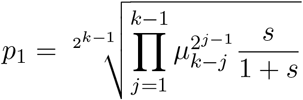

is independent of the *δ*_*i*_, so relaxing selection only decreases the probability that an adapted individual eventually fixes.

## 2 Wright-Fisher Model

The above reasoning translates readily to the Wright-Fisher model. The latter is characterized by discrete, non-overlapping generations, where the pedigree of the next generation is determined by a fitness-weighted sampling of the previous generation and mutations that occur independently in each offspring. Where previously (§1.1) we approximated the numbers of individuals of rare types via a birth-and-death process, here, we use a Galton-Watson branching process [18] with a Poisson distributed offspring number. The latter allows us to once again write a recurrence equation for the probability, *p*_1_ that a given individual is ancestral to an adapted individual that carries *k* mutations and is destined to fix.

### 2.1 Poisson-Galton-Watson Approximation

We again assume a population of fixed size *N*. In this discrete-generation formulation we can implement fecundity selection by assuming that an individual carrying *i<k* mutations is “chosen” to be the parent of an individual in the next generation with weight *w*_*i*_ 1 (*N*.*δ*., carrying fewer than *k* mutations is deleterious). Thus, if the *j*^th^ individual (we assign an arbitrary ordering to the population) carries *i*_*j*_ mutations and has *X*_*j*_ offspring in the next generation, the joint distribution of the *X*_*j*_ is multinomial:

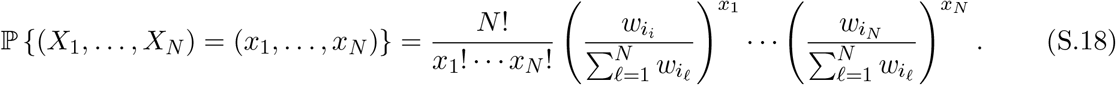

Using the grouping property of the multinomial distribution, *X*_*j*_ is binomially distributed:

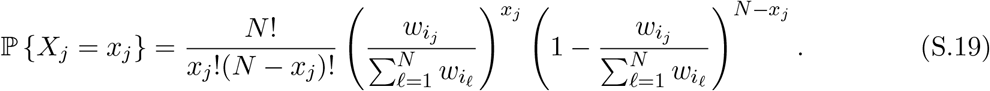

Now, letting 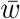 denote the population mean fitness,

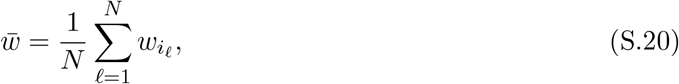

we have

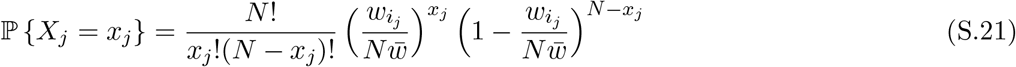

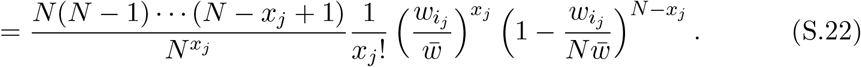

Taking the large population limit, we get

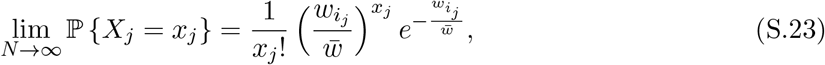

*i*.*e*., the *j*^th^ individual has a Poisson-distributed number of offspring with rate equal to their relative fitness, 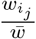. A similar calculation shows that for any arbitrarily chosen *k* individuals (*k* fixed and not growing with *N*), their offspring numbers will be approximately independent Poisson random variables with rates equal to the relative fitness of the individuals.

While this gives us a characterization of an individual’s reproductive output, unless the population is completely neutral, an individual’s relative fitness will depend on the state of the whole population. If, however, as before, we restrict our attention to the situation where the number of individuals carrying one or more mutations is negligible compared to *N*, then

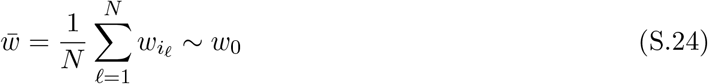

as *N* → 1 and thus, to first approximation, the *j*^th^ individual produces offspring with Poisson rate 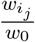 that is independent of the composition of the population. Thus, if *N* ≫ 1, we may approximate a small subpopulation of the Wright-Fisher process by a Galton-Watson branching process with Poisson distributed offspring.

Without loss of generality, we can take *w*_0_ = 1, provided we scale the *w*_*i*_ appropriately, taking *w*_*i*_ = 1 - *δ*_*i*_ for 0 *<i< k* and *w*_*k*_ = 1 + *s*. As previously, we can model additive selection by taking *δ*_*i*_ = *bi* and multiplicative selection by taking

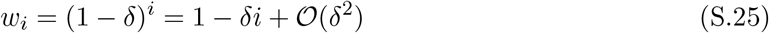

for 0 *<i< k*, for some *δ* ≪ 1.

### 2.2 Galton-Watson Recurrence

As before, we wish to compute *p*_*i*_, the probability that an individual carrying *i* mutations is ancestral to a mutation destined to fix. We will again do this recursively. We first compute the recurrence relation for an arbitrary Galton-Watson process. Then, in the next section, we will specialize to the Poisson-Galton-Watson process derived above.

No individual survives more than one generation, so an individual is ancestral to an individual destined to sweep if and only if one of its offspring is ancestral to the selected individual. As before, an offspring may or may not acquire an additional mutation, and we assume that the probability of multiple mutations is negligible (*µ*_*i,j*_ = *µ*_*i*_ if *j* = *i* + 1, and is zero otherwise). Thus, as previously 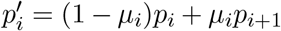 (S.3) is the probability that some given offspring of an individual carrying *i* mutations is ancestral.

To obtain the probability that at least one offspring is ancestral, we use the probability generating function for the offspring distribution: if *X* is the (random) number of offspring, then

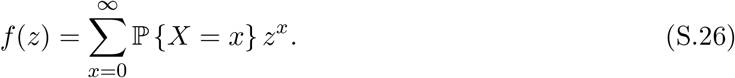

(for a Poisson distribution with rate .*\*, we have *f* (*z*) = *e*^*λ* (*z-*1)^). Conditional upon *X* = *x* the probability that *no* offspring is ancestral is 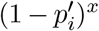, whence the unconditional probability that no offspring is ancestral is

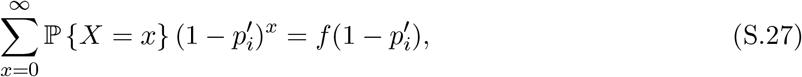

so that the probability that at least one is ancestral is 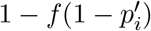, yielding the recurrence

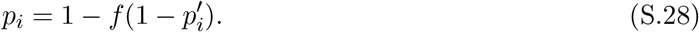

### 2.3. Solution in the Poisson Case

We now focus on the case of a Poisson-distributed number of offspring, corresponding to our approximation to the Wright-Fisher model (§2.1). In this case, (S.28) becomes

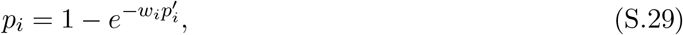

where 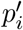 is given by (S.3), and as above, we adopt the convention that *w*_0_ = 1, so that each individual’s relative fitness is simply *w*_*i*_.

This can be solved exactly using the Lambert *W* -function [2]. *W* (*z*) is the multi-function satisfying the relation *W* (*z*)*e*^*W* (*z*)^ = *z*. Two branches of the *W* -function are real valued, *W*_0_(*z*) and *W*^−1^(*z*), which map [− *e*^−1^, ∞) to [−1,∞) and [−*e*^−1^, 0) to (−∞, −1], respectively. The two branches meet at the point (−*e*^−1^, 1)). These real-valued branches satisfy a one-sided inverse identity:

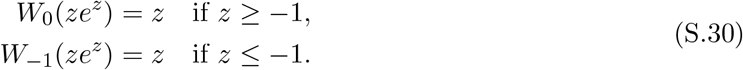

Moreover,

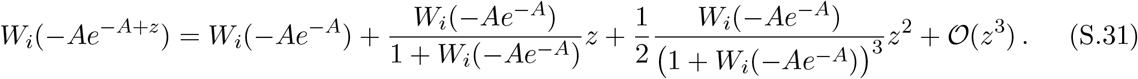

see [16, Equation (2.40)]. If *A* ≤ 1, using (S.30), this reduces to

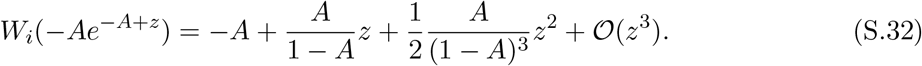

To use the *W* -function to solve the recurrence, we rearrange (S.29) as:

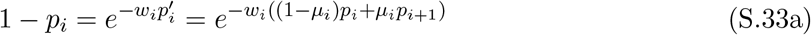

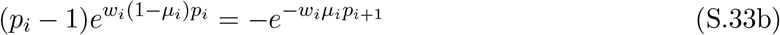

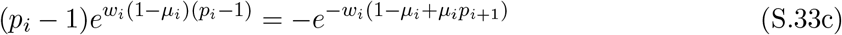

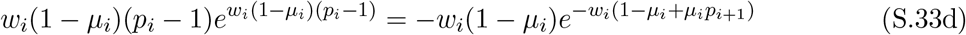

Now, 0 *< w*_*i*_ ≤ 1, 0 ≤ *p*_*i*_ ≤ 1, and 0 *< µ*_*i,i*+*l*_ ≪ 1, whence *w*_*i*_(1 − *µ*_*i*_)(*p*_*i*_ − 1) ≥ −1. Applying *W*_0_ to both sides of (S.33d), using (S.30) and (S.31) we conclude that

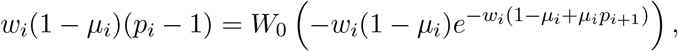

So

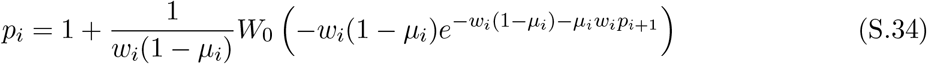

which we can iterate and simplify as above.

Using (S.32), we can approximate (S.34) as

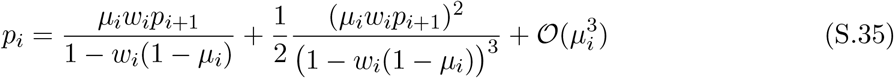

since −*w*_*i*_(1 − *µ*_*i*_) ≥ −1.

#### 2.3.1. Initial condition *p*_*k*_

To compute the initial value *p*_*k*_, we note that a selectively favoured individual fails to establish itself if and only if all of its offspring fail to establish. Let *p* denote the probability of successful establishment and let *q* = 1 − *p*. Letting *f* (*z*) be the probability generating function for the offspring distribution, as above (S.26), we must have *q* = *f* (*q*). In the Poisson case, this gives us *q* = *e*^*wk*(1*−q*)^. As before, we can solve this using the Lambert *W* -function. After rearranging, this is

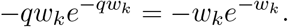

This always has the trivial solution *q* = 1, since − *qw*_*k*_ = − *w*_*k*_ ≤ − 1, but also a non-trivial solution with − *qw*_*k*_ =≥− 1 (*cf*. (S.30)):

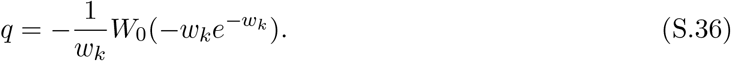

(recall *w*_*k*_ = 1 + *s* for *s>* 0).

Assuming *s* ≪ 1, we can find an elementary approximation to the Lambert *W* -function: let ε (*z*) = *ze* ^*z*^, so *W* (*z*) is ε^−1^ (*z*). Then, the *n* ^th^ derivative of ε (*z*) is

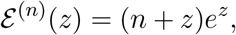

so, the second order Taylor polynomial for ε (*z*) at *z* = −1 is

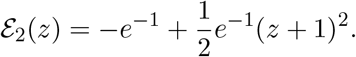

We can invert ε_2_(*z*) using the quadratic formula: setting *y* = ε_2_(*z*) and expanding, we see that

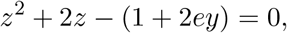

whence

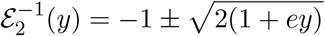

and

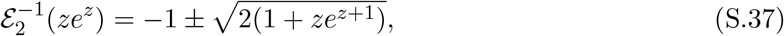

the two roots corresponding to the two real branches of the Lambert *W* function: + gives the *i* = 0 branch, which takes value in [−1, ∞), whereas - gives the *i* = -1 branch, which takes values in (−1, ∞].

Applying this approximation using *z* = −*w*_*k*_ = −(1 + *s*), we find

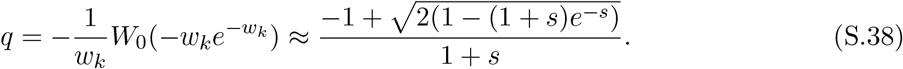

Expanding the numerator using the binomial series, we find 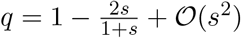, whence

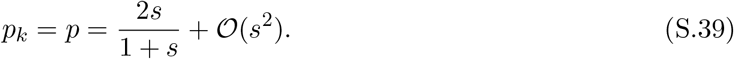

*N*.*δ*., this approximation to the fixation probability of an allele with selective advantage *s* was first obtained in [8], without reference to the exact solution via the Lambert *W* -function. The factor of two difference between the Wright-Fisher (S.39) and Moran models (S.6) is a consequence of a similar factor of two in the effective population sizes [7, §3.7].

Iterating (S.35) starting from 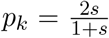 and assuming as previously, that *µ*_*i*_ ≪ *δ*_*i*_ ≪ 1, we get that to lowest order,

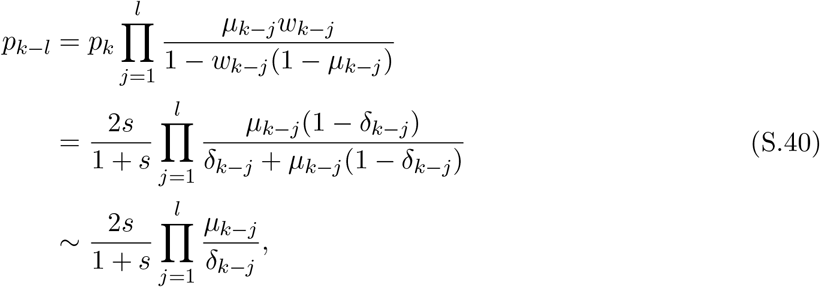

which corresponds to the case when *i* = 1 in the Moran model (S.11), (S.12). Interestingly, unlike the Moran model, there are no other regimes that need to be considered. As previously, the waiting time to the first ancestor is 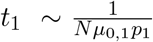, and our argument in §1.3 shows that relaxing selection decreases the mean time to adaptation in the Wright-Fisher model as well.

## 3 Justifying the Assumption that Mutants are Rare

Our arguments have been based upon the assumption that mutant individuals are rare, and that their numbers are negligible as compared to the number of wild-type individuals. We argued that this was a consequence of negative selection. Here, we briefly discuss some results on the maximum population size of birth-and-death and branching processes that show that provide evidence for that claim and quantify the strength of selection necessary. A rigorous proof would require additional coupling arguments to formalize the comparison between the Moran and Wright-Fisher models and the corresponding approximations; we omit these in the interest of brevity.

### 3.1 Moran case

We will consider the total progeny of a given 1-mutant. To simplify our presentation, in what follows we will measure time from the birth of that individual. We let *X*(*t*) denote the total progeny of our focal individual alive at time *t* (so *X*(0) = 1) and let

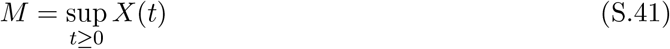

be the maximum number of progeny alive at any given time. Here, we present an estimate of the cumulative distribution function (c.d.f.) of *M*, ℙ{*M* ≥ *m*}.

We start with a simple observation: if *T* = inf {*t* : *X*(*t*) ∈{0, *m*}} then *M* ≥ *m* if and only if *X*(*T*) = *m*. If *X*(*t*) is a birth-and-death process with birth and death rates *δ* and *d* respectively, then this probability is well known (see *e*.*g*., [13], [9, §3.7], or [5, Theorem 6.1]):

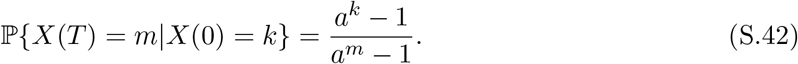

where 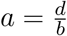.

We cannot directly apply this to our process, where individuals have different birth rates, *δ*_*i*_ = 1 − *δ*_*i*_, according to the number of mutations that they carry. We can, however, use it to obtain an upper bound: all individuals have birth rate no greater than *δ*_1_ = 1 − *δ*_1_ and death rate one, and thus will be fewer in number than the birth-and-death process, *X*(*t*), with those rates and *X*(0) = 1. For this *X*(*t*),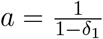,so

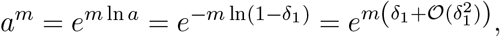

and we see that the probability that *M* ≥ *m*,

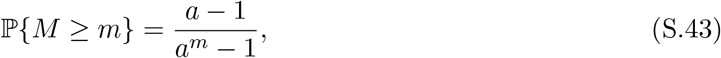

is exponentially small provided 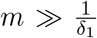. Thus, to prevent the number of descendants from ever reaching size 𝒪 (*N*), we need that 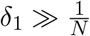. Note that

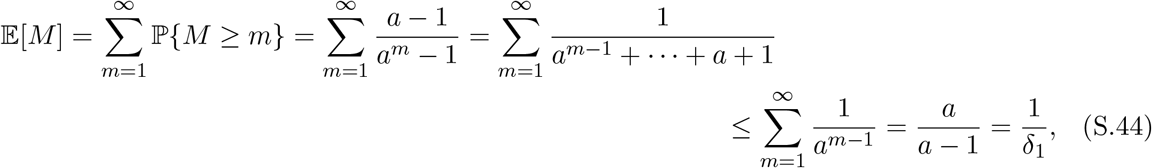

with approximate equality when *a* ≫ 1 (*δ*_1_ ≪ 1). In particular, for very small *δ*_1_ and large *µ*_0_—leading to a number of overlapping lineages—the rare mutant approximation can fail.

### 3.2 Wright-Fisher case

The argument for the Wright-Fisher case is almost identical to §3.1. We will bound the total number of mutant progeny of a single individual by a Poisson-Galton-Watson branching process *X*(*t*) where all individuals have fitness *w*_1_ = 1 − *δ*_1_, and *X*(0) = 1.

Define *M* and *T* as previously (where now time is measured in generations). The result (S.42) continues to hold for the Galton-Watson process (see [12, §3]) provided there exists *a>* 1 such that *f* (*a*) = *a*, where *f* (*z*) is the p.g.f. for the offspring distribution (see (S.26) above) of an individual with one mutation (hence *m* = *f*^′^ (1) *<* 1).

Just as we used the Lambert *W* function to compute the extinction probability *q* for the beneficial mutation, we can use it to compute *a* for a Poisson(*w*_1_) offspring distribution. Recall that for this distribution, the p.g.f. is *f* (*z*) = *e*^*w1*(*z*𢄡)^, so *a* = *f* (*a*) if and only if

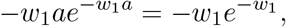

which has two solutions, *a* = 1 and

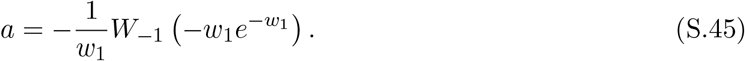

In particular, if *w*_1_ = 1 − *ϵ*for some small *ϵ*, then using (S.37) above, we have

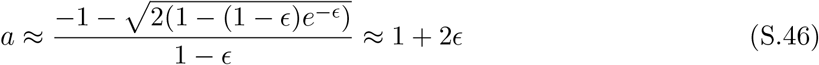

Now suppose *ϵ*= *ϵ*_*N*_ → 0 as *N* →1. Then,

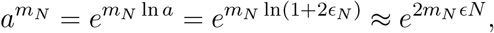

so the probability that *M* ≥ *m*_*N*_ tends to 0 as *N* → 1 if 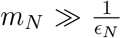, and to 1 if 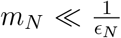, and to a non-zero constant if 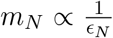. In particular, the number of deleterious mutants could reach 𝒪 (*N*) if 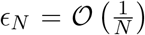, *i*.*e*., if selection is weak. This gives us the lower bound on the strength of selection as discussed above.

### 3.3 Conditioning on Not Fixing is Equivalent to Assuming Negative Selection

The above arguments bound the number of *i*-mutants, *i< k*, for which the mutations are deleterious. We must also, however, bound the number of *k*-mutants that do not go on to fixation, which, due to epistasis, have selective advantage *s*. These lines will eventually go on to extinction, but we must ensure they remain small.

This, fortunately, follows almost immediately from our previous work. *Conditioned upon extinction*, a birth-and-death process with birth rate *δ* and death rate *d, b> d*, is equal in law (*i*.*e*., has the same probability distribution) as a birth-and-death process with birth rate *d* and death rate *δ* (see [15, Lemma B.2]). Applying this to lineages of *k*-mutants conditioned on extinction, we have *δ* = 1 + *s, d* = 1, and we can take 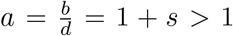 in (S.43) to estimate the maximum number of *k*-mutants.

Similarly, *conditioned upon extinction*, a Galton-Watson process with p.g.f *f* (*z*) for the offspring distribution and extinction probability *q <* 1 (recall *f* (*q*) = *q*) is equivalent to a Galton-Watson process with offspring p.g.f. 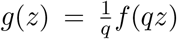 (see *e*.*g*., [1, Theorem I.8.1]). Thus, *g*(*a*) = *a* for 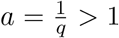. Applying this to the *k* mutants, we have 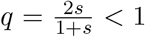 so taking 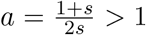 in (S.43) gives us a bound on their maximum number.

## 4 Bounding the Adaptation Time

We now turn our attention to estimating the expected waiting time from the birth of the ultimate ancestor carrying 1 mutation to the birth of the adapted individual. There is a direct line of descent between the 1-ancestor and the adapted individual. In what follows, we will refer to individuals on this line as ***ancestral***, and those carrying *i* mutations as *i*-***ancestors***. Thus an *i* ancestor gives birth at rate *i*, and its offspring carries *i* + 1 mutations with probability *µ*_*i*_.

Imagine following the genealogical tree between the ultimate ancestor and adapted individual: we follow the branch corresponding to each ancestral individual until they give birth to their successor in the line of descent, after which we follow the successor’s branch. Over it’s lifespan, an *i*-ancestor gives births at the times of a Poisson process with rate *δ*_*i*_, but most of these will not be ancestral: the probability that a given offspring is ancestral is 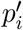 (S.3). By the thinning property of the Poisson process (see *e*.*g*., [3, §3.1]), the *i*-ancestor gives birth to would-be ancestors (only the first will be truly ancestral) as a Poisson process with rate 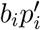 *i*.*e*., the waiting time birth of next ancestor is exponentially distributed with rate 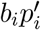. Again, the probability that one of these ancestors is an *i* + 1-ancestor is *µ*_*i*_, further thinning the process, so that the waiting time from the first *i*-ancestor to the first *i* + 1 ancestor is exponentially distributed with rate 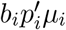.

The total adaptation time is the sum of these exponential waiting times, but we must *condition upon the fact that the adapted individual appears before the line of descent goes extinct*. This is a difficult, and possibly intractable problem. Here, we will content ourselves with a coarse approximation: we will simplify the processes by *underestimating* the number of mutants, which we do by considering a birth-and-death process *X*(*t*) with death rate 1 and birth rate *δ* = *δ*_*k−*1_ = 1 − *δ*_*k−*1_ and *X*(0) = 1 (compare §3.1), and we will assume that all mutation rates are equal *µ*_*i*_ *≡µ*. We then have 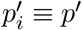.

The *uncondtioned* waiting time to the adapted individual, say *T*_a_, is then a sum of *k* exponentially distributed random variables with rate. λ = *bp*^′^*µ, i*.*e*., an Erlang or Gamma distributed random variable with probability density function (p.d.f)^1^

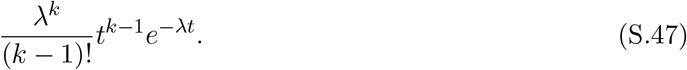

The total line of descent, on the other hand, has lifespan *L* with cumulative distribution function (c.d.f.) [11, §3]

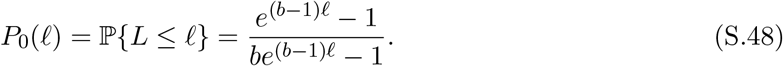

*N*.*δ*., *b<* 1, so *P*_0_(∞) = 1.

Now, suppose *L* = 𝓁. The density of *T*_a_ conditional upon *T*_a_ ≤ 𝓁 is

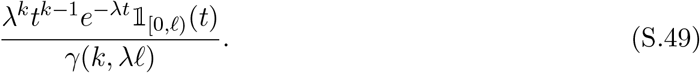

where 𝟙_[0,𝓁)_(*t*) is the indicator function of the interval [0, *L*)^2^ and *γ* (*a, z*) is the ***incomplete gamma function*** [4, §8.2.1]:

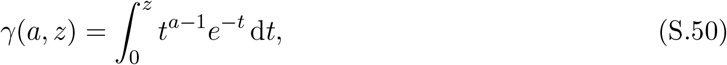

the latter arising as

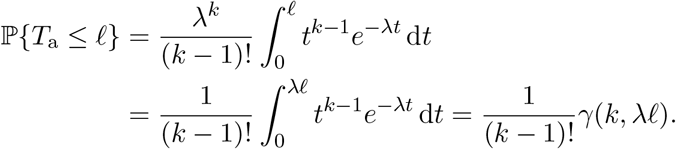

Integrating (S.49) against 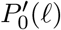, (S.48), gives us the p.d.f. of *T*_a_ conditional on the random lifespan *L* (as opposed to the fixed value *L* = 𝓁):

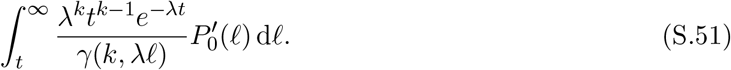

We can use this to estimate the expected adaptation time, 𝔼 [*T*_a_]:

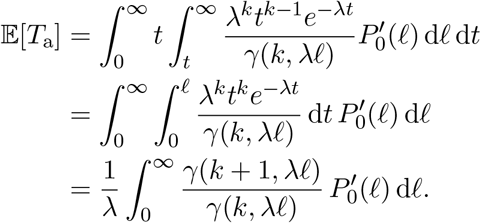

This latter integral does not appear to be analytically tractable, but we can obtain a suitable bound by noting that for integer values of *k*, [4, §8.4.7],

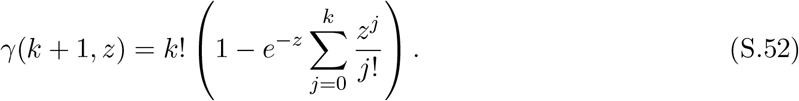

In particular,

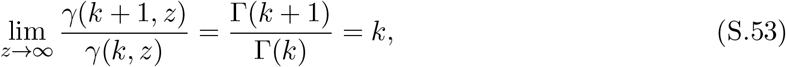

and 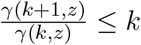 for all *z*, whereas for *z* ≤ *k* + 1,

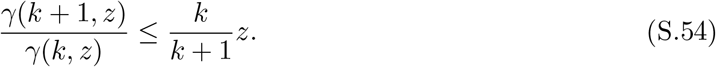

Thus,

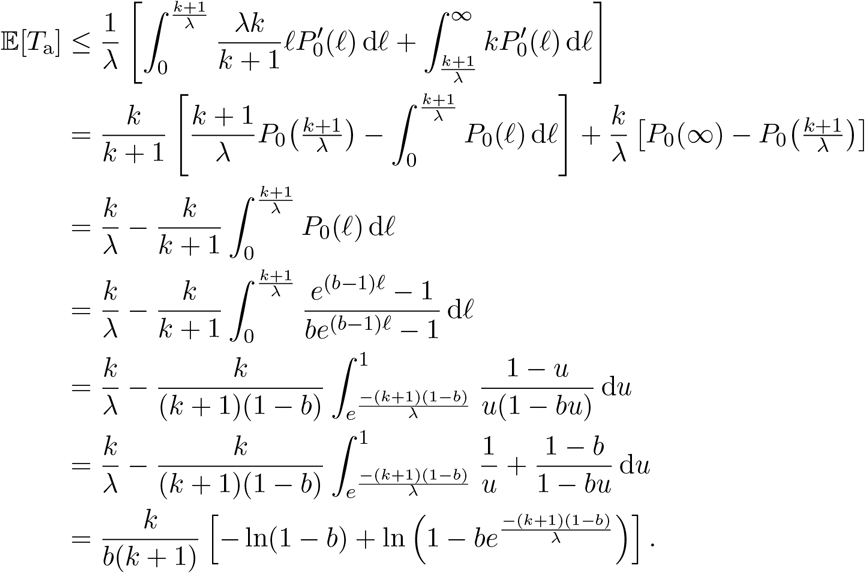

Recalling that *b* = 1 − *δ*_*k-*1_ and. *λ* = *δ*^′^*µ* ≪ 1, the latter is approximately

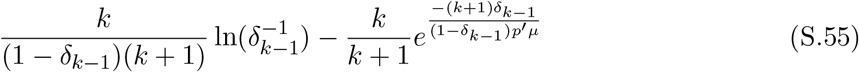

where the latter term is negligible compared to the first.

Finally, it is a standard result (see *e*.*g*., [5, Theorem 6.3]) that the fixation time for an allele with selection coefficient *s* in a population size *n* is approximately 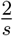 ln *n*.

## Footnotes

1 Informally, a random variable *X* has p.d.f. *f* (*x*) if ℙ*{X*∈*2* [*x, x* + d*x*)*}* = *f* (*x*) d*x*.

2 𝟙_[0,𝓁*`*)_(*t*) = 1 if *t* ∈ [0, 𝓁), and is zero otherwise.

